# Dppa2/4 promotes zygotic genome activation by binding to GC-rich region in signaling pathways

**DOI:** 10.1101/2020.03.18.998013

**Authors:** Hanshuang Li, Chunshen Long, Jinzhu Xiang, Pengfei Liang, Yongchun Zuo

**Affiliations:** State Key Laboratory of Reproductive Regulation and Breeding of Grassland Livestock, College of Life Sciences, Inner Mongolia University, Hohhot 010070, China

**Author notes:** These authors contributed equally to this work as first authors (Hanshuang Li, Chunshen Long).

**Keywords:** Dppa2/4, zygotic genome activation, GC-rich region, decisive signaling pathways, Alppl2

## Abstract

Developmental pluripotency associated 2 (Dppa2) and Dppa4 as positive drivers were helpful for transcriptional regulation of ZGA. Here, we systematically assessed the cooperative interplay between Dppa2 and Dppa4 in regulating cell pluripotency of three cell types and found that simultaneous overexpression of Dppa2/4 can make induced pluripotent stem cells closer to embryonic stem cells. Compared with other pluripotency transcription factors (TFs), Dppa2/4 tends to bind on GC-rich region of proximal promoter (0-500bp). Moreover, there was more potent effect of Dppa2/4 regulation on signaling pathways than other TFs, in which 75% and 85% signaling pathways were significantly activated by Dppa2 and Dppa4, respectively. Notably, Dppa2/4 also can dramatically trigger the decisive signaling pathways for facilitating ZGA, including Hippo, MAPK and TGF-beta signaling pathways and so on. At last, we found that Alkaline phosphatase placental-like 2 (Alppl2) was significantly activated at the 2-cell stage in mouse embryos and 4-8 cell stage in human embryos, further predicted that Alppl2 was directly regulated by Dppa2/4 as a candidate driver of ZGA to regulate pre-embryonic development.

## Introduction

Zygotic genome activation (ZGA) as the first key concerted molecular event of embryos, which major wave takes place in two-cell stage in mouse embryos is crucial for development (Hu et al, 2019; in; Vastenhouw et al, 2019). During the process of ZGA, epigenetically distinct parental genomes can be reprogrammed into totipotent status under the coordinated regulation of pivotal factors (Hu et al, 2020; Li et al, 2020). Nowadays, several ZGA decisive triggers are being well identified, such as the TF Dux, expressed exclusively in the minor wave of ZGA, was discovered to further bind and activate many downstream ZGA transcripts (De Iaco et al, 2017; Hendrickson et al, 2017; Whiddon et al, 2017). The endogenous retrovirus MERVL was also defined as a 2-cell marker (Macfarlan et al, 2012; Yan et al, 2019). And Zscan4c was newly reported as important inducers of endogenous retrovirus MERVL and cleavage embryo genes (Zhang et al, 2019). However, we still know little about how TFs drives complex transcriptional regulation events of ZGA and the coordination among them. For example, Dux is discovered to have only a minor effect on ZGA in vivo, whereas is essential for the entry of embryonic stem cells (ESCs) into the 2C-like state (Chen & Zhang, 2019).

Developmental pluripotency associated factor 2 (Dppa2) and developmental pluripotency associated factor 4 (Dppa4) as small putative DNA-binding proteins were observed expressed in pluripotent cells and the germline (Kang et al, 2015; Madan et al, 2009; Maldonado-Saldivia et al, 2007). Both of these two proteins have an N-terminal conserved SAP (SAF-A/B, Acinus, and PIAS) motif, which is important for DNA binding, and a C-terminal associated with domain histone binding (Aravind & Koonin, 2000; Maldonado-Saldivia et al, 2007; Masaki et al, 2010). Embryonic development is impaired when Dppa2 and 4 are depleted from murine oocytes, suggesting that the protein plays an important role in the early pre-implantation period (Madan et al, 2009). Knockdown of either Dppa2 or Dppa4 reduces the expression of ZGA transcripts (Eckersley-Maslin et al, 2019), and these two factors also act as inducers of Dux and LINE-1 transcription in mouse embryonic stem cells (mESCs) to promote the establishment of a 2C-like state (De Iaco et al, 2019), confirming that these two proteins are necessary to activate expression of ZGA transcripts.

Additionally, both Dppa2 and Dppa4 associate with transcriptionally active chromatin in mESCs (Engelen et al, 2015; Masaki et al, 2007), which function as key components of the chromatin remodeling network that governs the transition to pluripotency (Hernandez et al, 2018; Masaki et al, 2007). Other studies also have proved that these two proteins have pluripotency-specific expression pattern (Chakravarthy et al, 2008) and can be used as pluripotency markers to recognize induced pluripotent stem cells (IPSCs) (Kang et al, 2015; Klein et al, 2018). Understanding how Dppa2 and Dppa4 function would provide a better understanding of the mechanisms these two factors in transcriptional regulation.

Here, we described a genome-wide dynamic binding profile of Dppa2 and Dppa4 among different cell types: mouse embryonic fibroblasts (MEFs), IPSCs and ESCs (under Dppa2/4 single- and double-overexpressed treatment). The chromosome and sequence preference of Dppa2 and 4 binding distributions were further explored. Compared with other TFs, Dppa2 and 4 tend to binding on GC-rich region of proximal promoter to activating majorities of signaling pathways. Next, we identified the unique and common target genes of Dppa2 and 4 in ESCs, in which there was more significant effect of Dppa4-bound on genes related to ESCs than Dppa2. Notably, Dppa2/4 can also comprehensively activate some signaling pathways associated with developmental reprogramming for facilitating ZGA. At last, we predicted Alppl2 that directly activated by Dppa2 and Dppa4, which may as a new activator acts in folate biosynthesis, promoting the progress of ZGA. Taken together, deciphering the transcriptional regulation events of key factors during ZGA is crucial to elucidate the potential mechanism of early embryo development.

## Results

### The cooperative interplay between Dppa2 and Dppa4 in regulating cell pluripotency

To systematically investigate the target binding effects of Dppa2 and Dppa4, principal component analysis (PCA) was performed to validate the binding impacts of these two TFs in MEFs, IPSCs and ESCs. We found that Dppa2 and Dppa4 established a dynamic modification roadmap among different cell types (Fig 1A), in which, simultaneous overexpression of Dppa2 and 4 can make IPSCs closer to the ESCs state and the same result also can be confirmed in clustering analysis, indicating that Dppa2/4 can further facilitate the strengthening of cell pluripotency, especially significant is cooperative effect of both (Fig 1A). Next, we comparatively observed the binding preference of Dppa2 and Dppa4 on different chromosomes in three cell types, the overall distribution of Dppa2 and Dppa4 binding peaks in ESCs was higher than in other two cell types. Among them, Dppa2 and 4 are inclined to bind on chromosomes 2, 4, 5, 8, 11, 15 and 17 (Chr2, Chr4, Chr5, Chr8, Chr11, Chr15, and Chr17), especially Chr4 and Chr5, which are the major target chromosomes (Appendix Fig S1A). However, relatively few peaks are located on the sex chromosomes. The results showed that the binding of Dppa2/ 4 had not only cell type specificity, but also chromosome preference.

**Figure 1.**
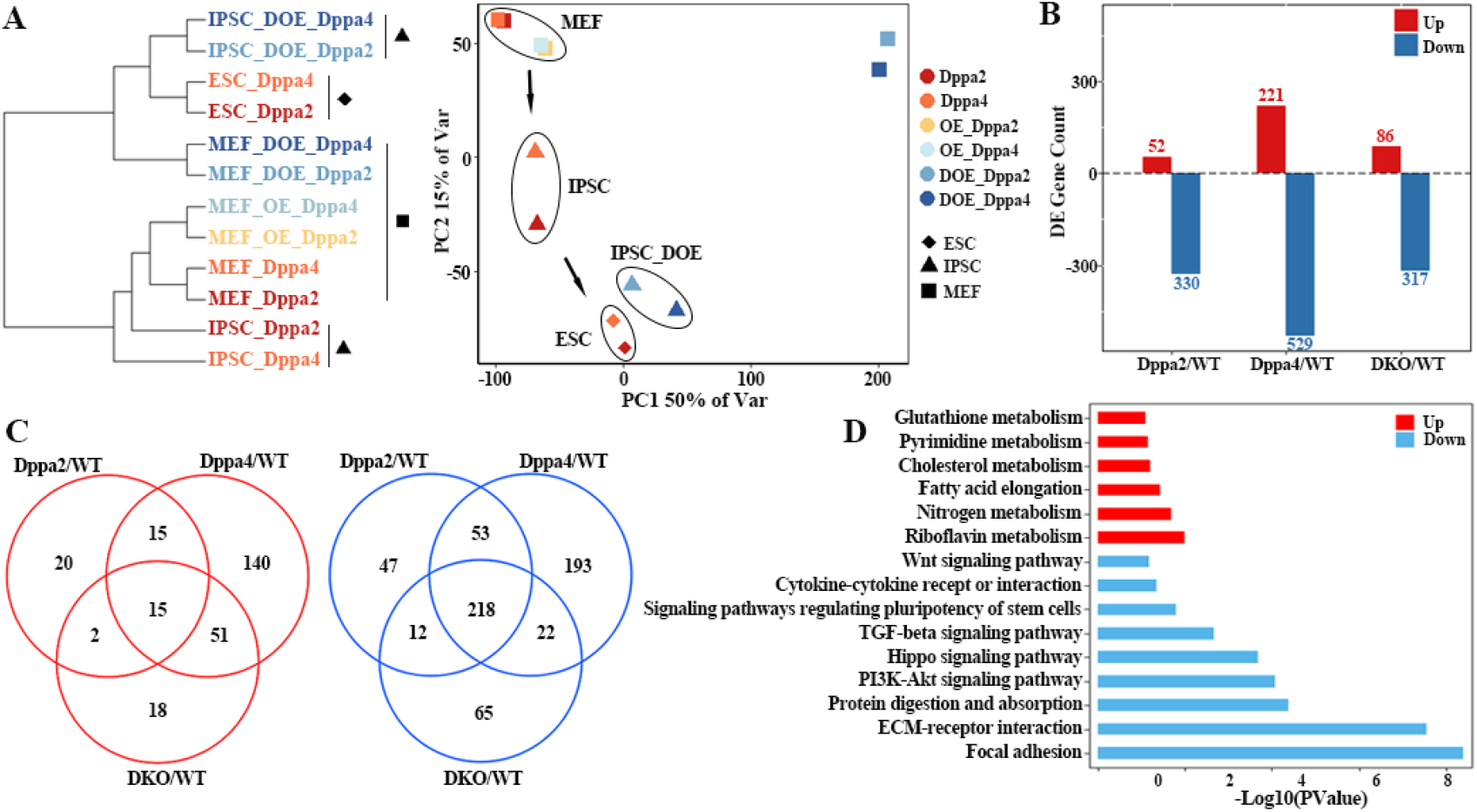
Dppa2 and Dppa4 synergistically regulate cellular pluripotency. A Hierarchical clustering and principal component analysis according to the target binding patterns of Dppa2 and 4 in Dppa2/Dppa4 single-, double-overexpression MEFs, IPSCs and ESCs. The coordinates indicate the percentage of variance explained by each principal component. B Analysis of differentially expressed genes (DEGs) between Dppa2/Dppa4 single-, double-knockout treatment (Dppa2-KO, Dppa4-KO, DKO) and control WT ESCs. C Overlap among DEGs following Dppa2/Dppa4 single- and double-knockout treatment compared with control WT ESCs. The left and right Venn diagram representing the overlap of up-regulated and down-regulated genes, respectively. D GO pathway analysis of up-regulated and down-regulated overlap genes, the up- and down-regulated genes enrichment pathways are shown in red and blue, respectively.

To better understand the roles of Dppa2 and Dppa4 in gene regulation in ESCs, we performed differential expression analysis between Dppa2, Dppa4 single-, double-knockout treatment (Dppa2-KO, Dppa4-KO and DKO) and control (WT) ESCs (Fig 1B). Comparing with Dppa2 knockout, Dppa4 knockout resulted more ESC-related genes differential expression, of which the up-regulated gene was 221 and down-regulated genes were 529 (Fig 1B). Additionally, there were 218 down-regulated genes shared in Dppa2, Dppa4 single- and double-knocked ESCs (Fig 1C), which were enriched in multiple KEGG pathways related to pluripotency regulation and development, such as Wnt, Hippo PI3K-Akt signaling pathways, etc. (Fig 1D), likely reflect the similar influence of these two factors in the identity maintenance of ESCs. Interestingly, Dppa4-KO also affected the differential expression of a substantial number of unique genes, which may indicate the more important roles of Dppa4 in ESCs. Moreover, there were 15 differentially up-regulated genes shared in the Dppa2, Dppa4 single-, double-knockout treatment ESCs, these genes were mainly involved in Fatty acid elongation, Riboflavin metabolism, Nitrogen metabolism and other metabolism related pathways (Fig 1D), confirming that the synergistic regulation of these two factors in basic metabolism functions.

### Dppa2 and Dppa4 are prior binding to CG-rich region of proximal promoter

To examine whether there was bias in the genomic locations of Dppa2 and Dppa4-binding sites, we analyzed the genomic distributions of Dppa2 and Dppa4 peaks in MEFs, IPSCs and ESCs (Fig 2A). Among these three cell types, there were 20000 binding sites of Dppa2/4 on genes promoter regions. Approximately 40% peaks binding on Genebody, in which 1st exon and 1st intron are the prior binding genomic regions. While in contrast on enhancer regions, only 10% binding peaks of the two TFs (Fig 2A). By analyzing the distribution of Dppa2/4 binding sites with respect to the genes transcriptional start sites (TSS) in each cell type (Fig 2B), we found a major binding wave of Dppa2/4 within 5 kb of the TSSs upstream (0-5kb), while >15% of Dppa2/4 binding sites were located more than 5 kb of the TSSs, which formed a minor binding wave in the downstream of TSSs (−5 — -50kb) (Fig 2B). Moreover, there was a tendency for Dppa2/4 to bind on upstream TSSs in MEF, whereas in ESCs the minor binding wave was more significant. Next, we attempted precisely describe the sequence characteristics corresponding to the two binding waves, and found that the CG content of Dppa2/4 in these two waves is greater than 60% (Fig 2C). Furthermore, the percentage of CG contents of Dppa2, Dppa4 binding peaks are higher than in the peaks from other TFs (Fig 2D), suggesting that the two TFs preferentially bind in CG-rich regions (De Iaco et al, 2019).

**Figure 2.**
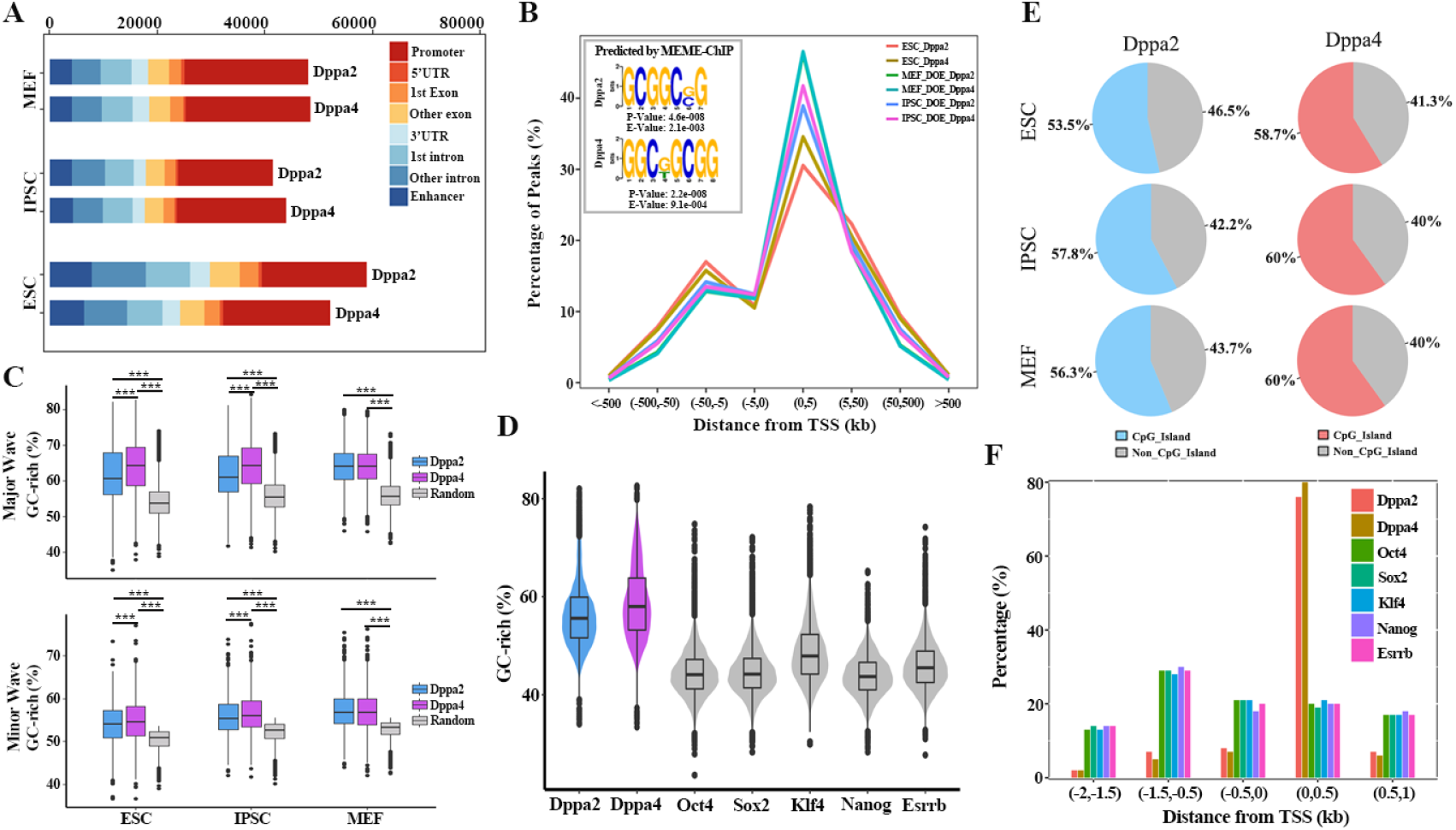
Dppa2 and Dppa4 tend to binding to CG-rich region. A The genomic distributions of Dppa2/4 binding peaks in Dppa2/4 OE MEFs, IPSCs and ESCs. B Distribution of Dppa2/ Dppa4-binding sites with respect to the TSS in ESCs, MEFs and IPSCs is shown. And Dppa2/Dppa4-recognized DNA motifs identified by MEME-ChIP in ESCs. MEME Suite was used to determine E value of motif. C Boxplot representing the percentage of CG nucleotide contents in the peaks from Dppa2, Dppa4 ChIP, and in a random shuffle of peaks. Upper and lower boxplots involved the peaks binding on the major (upstream 0 kb to 5 kb of TSS) and minor binding waves (−50,−5kb), respectively. D The percentage of CG nucleotide contents in the binding peaks from Dppa2, Dppa4 and other TFs in ESCs (data reanalyzed from (Chronis et al, 2017)). E Pie charts summarizing the percentage of CpG Island of the Dppa2, Dppa4 peaks binding on promoter in indicated three cell types (left row). F The distribution of Dppa2, Dppa4 and other TFs-binding sites with respect to the TSS in ESCs.

We further verified the hypothesis that the CpG Island is the targeted binding region of Dppa2/4, and found that about 60% peaks of Dppa2 and Dppa4 bind on promoter contained CpG Island in ESCs, of which the content CpG Island of Dppa4-binding is higher than Dppa2 (Fig 2E). Similar results were also observed in the other two cell types. Approximately 75% of the target gene contained CG-Island was also bound by Dppa2 and 4 in promoter regions, which is consistent with previous investigations that CpG Island primarily contained in the genes promoter regions (Appendix Fig S2A) (Morgan & Marioni, 2018). Interestingly, Dppa2/4 tends to bind to the proximal promoter (0-500bp) more than other TFs (Fig 2F). Next, we identified the binding motifs of these two TFs based on MEME-ChIP (Machanick & Bailey, 2011) and found the binding motif of these two TFs (Dppa2, P: 4.6e-008; Dppa4, P: 2.2e-008) is strictly conserved in these three cell types (Fig 2B and Appendix Fig S2B). These results collectively indicate that Dppa2/4 function by binding to GC-rich region of proximal promoter.

### Target regulation ability of Dppa4 is superior to that of Dppa2 in ESCs

By investigating the binding characteristics of Dppa2 and Dppa4 in MEFs, IPSCs and ESCs, we found both the peaks and target genes of Dppa2/4 in ESC were significantly higher than in MEFs and IPSCs, and the targets containing RPKM were also higher than those of the other two cell types. Moreover, about 70% sites are bound to the target genes promoter, and above 60% of these target genes contain RPKM value (Fig 3A). Notably, the binding signals (logarithmic conversion of RPKM) distribution of Dppa2 and 4 is similar in ESCs and IPSCs, which was mainly distributed between −2.5 and 2.5. On the contrary, the signals distribution trends are −2.5— −5 and 2.5—7.5 in MEFs, suggesting that the targeted regulation of Dppa2/4 has cells type specificity (Fig 3B). Additionally, Dppa2/4 binding on their targets depends on the differential of gene regulatory elements in different cell types. In ESCs, Dppa2/4 was more likely to bind to targets with a small Exon Count (0-5) and longer 1stExon (500-1500bp), which is inconsistent in MEFs. While in IPSCs, they were tending to bind on target genes with longer 5’UTR (1000-3000bp) and 3’UTR (1000-3500bp) (Appendix Fig S2C).

**Figure 3.**
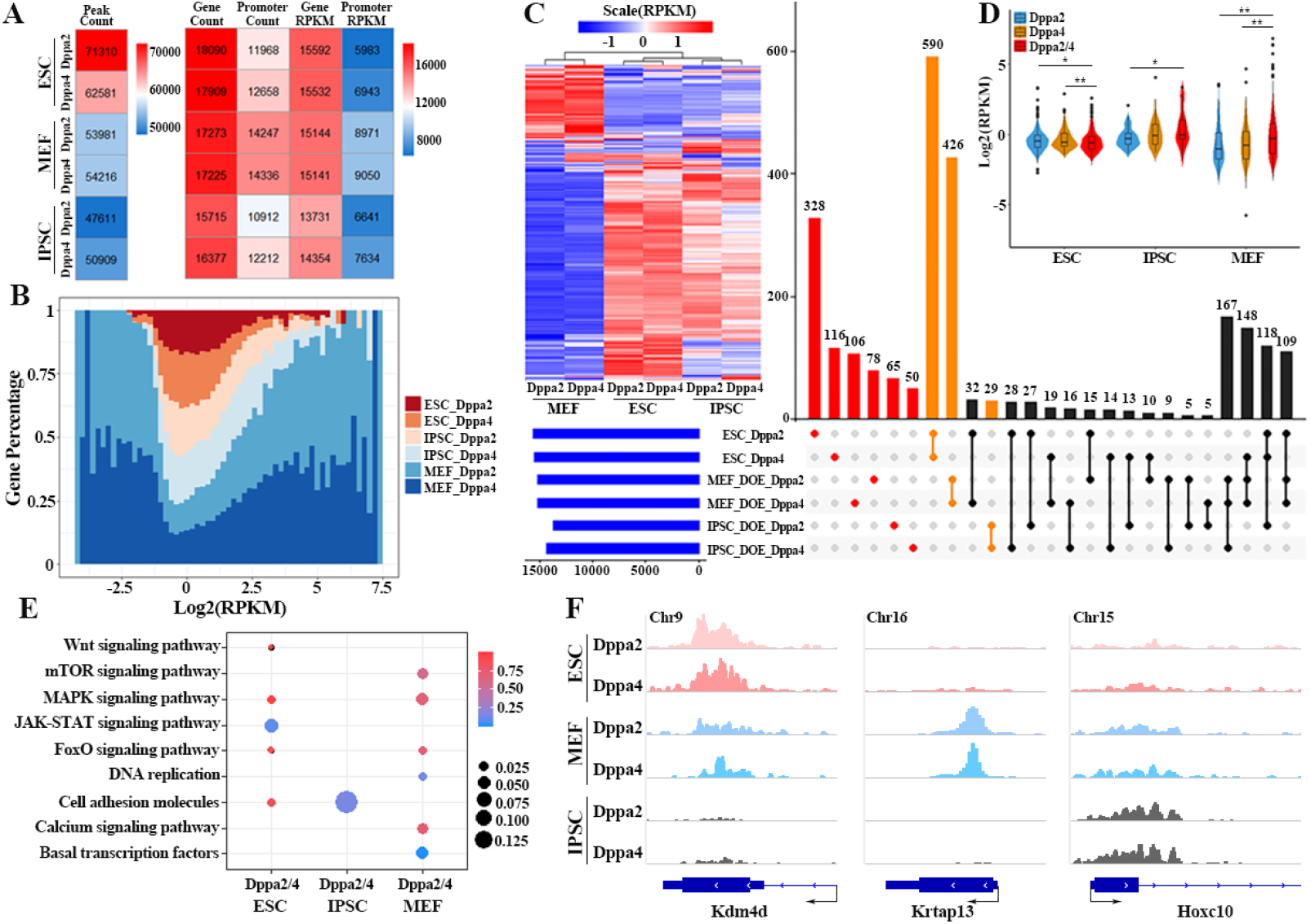
The target binding ability of Dppa2 and Dppa4 in MEFs, IPSCs and ESCs. A The heatmap showing the number of binding peaks, target genes, target genes on promoter, target genes containing RPKM of Dppa2, Dppa4 in MEFs, IPSCs and ESCs. B The density distribution of signal (RPKM) of Dppa2/4 binding on the promoter in MEFs, IPSCs and ESCs. And the x-coordinate represents the logarithmic transformation of RPKM. C Upset chart showing the target genes of Dppa2 and Dppa4 in three cell types (horizontal bar). The specific number of identified genes shared between different sets is indicated in the top bar chart corresponding to the solid points below the bar chart. Figure generated using Upset R package. Heatmap shows the binding signals on the target genes shared in Dppa2, Dppa4 and unique on each cell type. D The binding signals of unique and common target genes of Dppa2 and Dppa4 in the three cell types (the unique and common genes were involved in Figure 3C red and yellow bar, respectively). Differences are statistically significant. (*) P-value < 0.05; (**) P-value < 0.01; (***) P-value < 0.001, t-test. E KEGG pathway analysis of cell types unique genes co-regulated by Dppa2/4 (the genes were involved in Figure 3C yellow bar). F Genome browser view showing the Dppa2/4 binding signals on the representative genes of three cell types.

Next, we explored the characteristic of the target genes of Dppa2 and 4 in three cell types. There were 590, 426, 29 common target genes of Dppa2 and 4 in ESCs, MEFs and IPSCs, respectively, and the binding signals of these two TFs on these genes are shown in the heatmap (Fig 3C). Moreover, the binding signals of Dppa2/4 in ESCs and IPSCs more strongly than in MEFs. Especially, the coordinately binding ability of Dppa2/4 on genes were superior to individual regulation, in which Dppa4 was more potent (Fig 3D), which is different from previous study (Yan et al, 2019). The common target genes of Dppa2 and 4 in IPSCs are mainly involved in Cell adhesion molecules signaling pathway and in ESCs are predominantly enriched in Wnt and JAK-STAT signaling pathways, while in MEFs, they significantly participate in mTOR, Calcium, DNA replication and Basal transcription factors signaling pathways. Interestingly, MAPK and FoxO signaling pathways are present in both ESCs and MEFs (Fig 3E). Finally, the binding signals of Dppa2 and 4 on Kdm4d, Krtap13 and Hoxc10 are shown in Figure 3F, which as representative genes in ESCs, MEFs and IPSCs described the cell-specific binding of Dppa2/4 (Fig 3F).

### Dppa2/4 activate the major wave of signaling pathways associated with developmental reprogramming

We focused principally on the ESC data and collected 48 signaling pathways and 2290 related genes from KEGG database (Appendix Table S1). Compared to other TFs (Oct4, Sox2, Nanog, Esrrb and Klf4), Dppa2/4 can activate the major wave of signaling pathways, which is opposite of Oct4 (Fig 4A and Appendix Fig S2D). Among these, there are 32 significant signaling pathways and 5 targets genes were shared across Dppa2, Dppa4 and Oct4 binding comparisons, respectively (Fig 4B). Although the majority of target genes detected in Dppa2, Dppa4 and Oct4 binding were different, most of the significantly enriched signaling pathways were the same (adjPvalue <= 0.05) (Fig 4B). We also found that Dppa2/4 activate these signaling pathways by binding to GC-rich region (Fig 4C).

**Figure 4.**
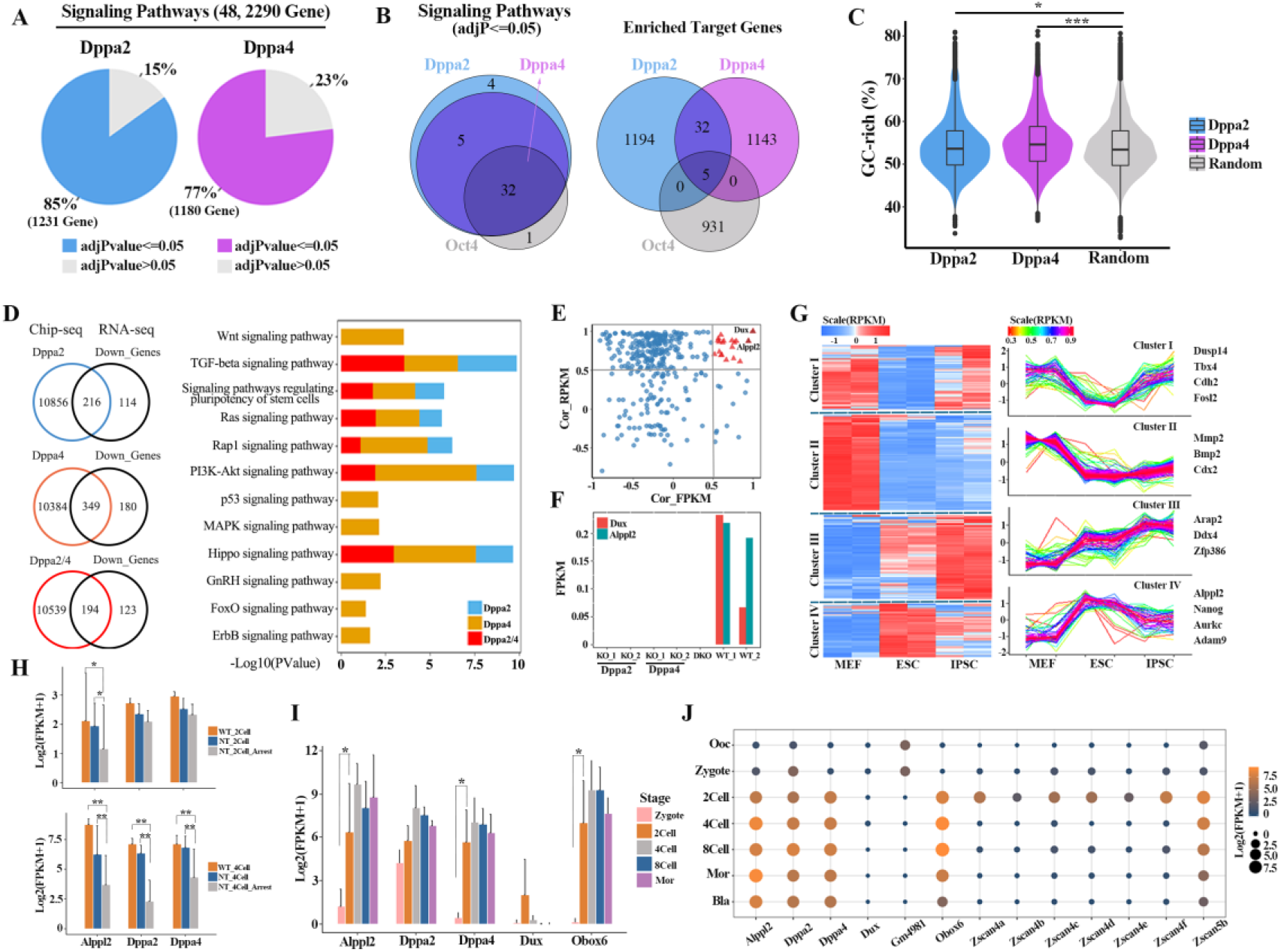
Dppa2/4 can significantly active majorities of the signaling pathways. A The significant enriched signaling pathways (adjPalve <= 0.5) and related target genes of Dppa2/4 in ESCs. B Significant enriched signaling pathways (adjPalve <= 0.5) and enriched target genes among Dppa2, Dppa4 and Oct4 in ESCs. C The percentage of CG nucleotide contents of Dppa2, Dppa4 binding on the targets in ESCs (the targets number was shown in Figure 4A). D Overlap of genes downregulated by Dppa2, Dppa4 single- and double-knockout treatment ESCs and Dppa2, Dppa4, Dppa2/4 ChIP-Seq peaks in ESCs. Significantly enriched KEGG pathways of these overlap genes, different groups involved the target genes bound by Dppa2, Dppa4 and Dppa2/4. E Scatter plot indicating the correlation between the target genes bound by Dppa2/4 (identified in figure 4D) and Dux, the horizontal axis represents the correlation of expression levels, and the abscissa represents the correlation of Dppa2/4 binding signals. F The expression patterns of Alppl2 and Dux in Dppa2/Dppa4 single-, double-knockout (Dppa2_KO, Dppa4_KO and DKO) and WT ESCs. G Fuzzy c-means (FCM) clustering analysis of target genes bound by Dppa2/4 (identified in figure 4A) in the indicated cell types. H The expression levels changes of Alppl2, Dppa2 and Dppa4 in NT, WT 2/4Cell embryos. Error bars represent average plus standard deviation of biological replicates (Mean+SD). Differences are statistically significant. (*) P-value < 0.05; (**) P-value < 0.01; (***) P-value < 0.001, t-test. I Dynamic changes in the expression patterns of representative genes during mouse embryos development. Differences are statistically significant. (*) P-value < 0.05; (**) P-value < 0.01; (***) P-value < 0.001, t-test. J The point plot shows the expression levels of corresponding genes in the development of mouse embryos, both the point size and color represent normalized gene expression levels (Log2(FPKM+1)).

Next, we sought to identify direct targets of Dppa2/4 by overlapping Dppa2 and Dppa4 ChIP-Seq peaks in promoters and genes down-regulated in response to Dppa2 and Dppa4 knockout. We found that more target genes regulated by Dppa4 than Dppa2 in ESCs, revealing Dppa4 may function as an ESC activator more prominent than Dppa2 (Fig 4D). Down-regulated genes that were directly bound by Dppa2/4 are enriched in majority of the signaling pathways related to the pluripotent maintenance and development, such as Signaling pathways regulating pluripotency of stem cells, TGF-beta, Ras, Rapl, PI3K-Akt and Hippo signaling pathways (Klein et al, 2018; Sasaki & Hiroshi). Among them, down-regulated genes that were directly bound by Dppa4 were unique enriched in KEGG pathways related to Wnt, p53, MAPK, GnRH, FoxO and ErbB signaling pathways (Fig 4D). Thus, several important signaling pathways related to development and reprograming were directly regulated by Dppa2/4, in which, Dppa4 has a greater effect on signaling pathway regulation than Dppa2. Previous findings have demonstrated that Dux acts as a downstream target gene for Dppa2/4 (Eckersley-Maslin et al, 2019; Yan et al, 2019), based on the strong association of binding signals and expression levels between these Dppa2/4 targets and Dux, a gene names Alkaline phosphatase, placental-like 2 (Alppl2) was screened out. Both the correlation of binding signals and expression levels between this gene and Dux were greater than 0.8 (Fig 4E). Intriguingly, this gene was completely silenced when Dppa2 and 4 single-or double-knockout in ESC, which is consistent with Dux (Fig 4F and Appendix Fig S3D). Therefore, it was demonstrated that Alppl2 may be a new direct downstream gene of Dppa2 and 4.

Fuzzy c-means (FCM) clustering analysis (Krinidis & Chatzis, 2010) of target genes bound by Dppa2/4 revealed that these genes can be categorized into 4 clusters (Fig 4G). Among them, cluster4 as an ESC-specific cluster contained Alppl2, further indicating that Alppl2 as an ESC-specific gene was directly regulated by Dppa2 and 4. To gain further insights into the roles of Alppl2, we reanalyzed the data of mouse SCNT embryos (Liu et al, 2016), and found that the expression level of Alppl2 in WT, NT normal 2/4-cell embryos is significantly higher than that in NT arrest embryos (Fig 4H). The results showed that Alppl2 plays a crucial role in promoting the success of SCNT reprogramming. Additionally, we found Alppl2 was significantly expressed at the 2-cell stage, and continued to Morula stage, consistent with the recent report that the specificity of ALPPL2 to naive pluripotency is conserved in mouse (Bi et al, 2020). Similarly, ALPPL2 was also observed to be up-regulated during the major ZGA (4-8 cell stage) in human embryos (Appendix Fig S3A and B) (data reanalyzed from (Xue et al, 2013)). Surprisingly, the expression level of Alppl2 was much higher than that of Dppa2 and 4 in the pre-implantation embryos of mouse and human (Fig 4I, Appendix Fig S3A and B). This result indicates that Alppl2 may be a new key activator of ZGA, which is directly bound by Dppa2 and 4, but whether a positive feedback loop of Alppl2 on Dppa2 and 4 has is unclear. Strikingly, Obox6 highly expressed at the 2-cell stage, which indicates another identity of Obox6 as ZGA key factor in addition to the reprogramming efficiency enhancer (Schiebinger et al, 2019) (Fig 4I and J, Appendix Fig S3C) (data from (Liu et al, 2018; Wang et al, 2018; Xue et al, 2013)). Furthermore, we also observed Zscan4 family members were mainly activated in the 2-cell phase, which is different from that the high expression of Gm4981 before ZGA (Fig 4G) (data from (Liu et al, 2016)), indicating that the importance of Zscan4 family genes in early mouse embryos development (Zhang et al, 2019).

### Alppl2 is predicted as a master driver participates in folate metabolism to regulate ZGA

Dppa2/4 act as epigenetic modifiers play an important role in the early pre-implantation period (Masaki et al, 2007; Nakamura et al, 2011). Apparently, Alppl2 were completely silenced when Dppa2 and 4 single-or double-knockout in ESC, implying the direct roles of these two factors in regulating Alppl2. Notably, we observed the enrichment motifs of Dppa2/4 binding on Alppl2 are conserved, which involved high CG content, implying that these two factors act on Alppl2 by a co-regulated way (Fig 5).

**Figure 5.**
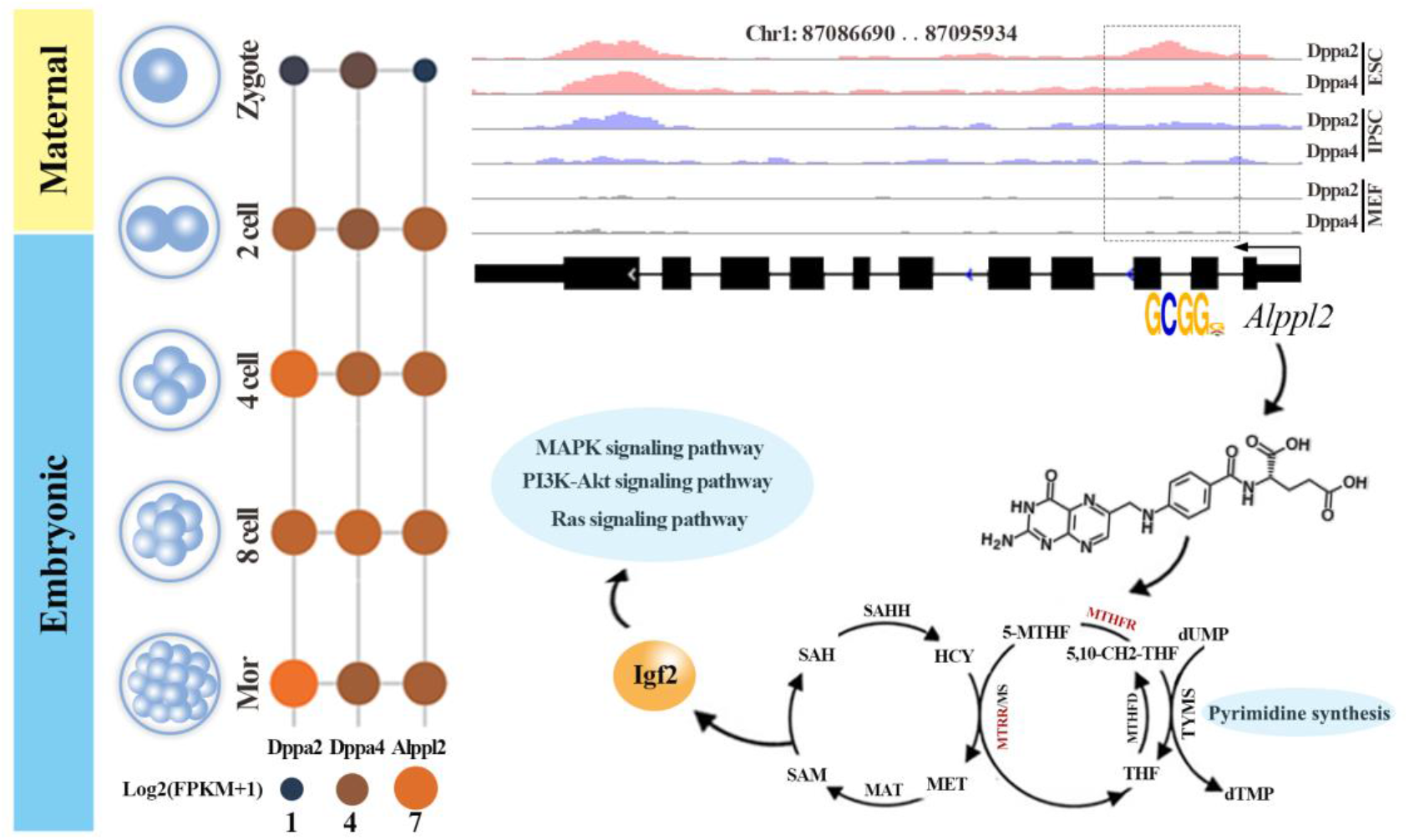
A predictive regulatory model of Dppa2/4 binding on Alppl2 for activating ZGA.

Previous reports have shown that Alppl2 is involved in the folate biosynthesis, and folate, which belongs to the family of B vitamins, plays essential roles in DNA synthesis, repair, and methylation (Geng et al, 2018; Panprathip et al, 2019). 5-methyltetrahydrofolate-homocysteine methyltransferase reductase (MTRR) and methylenetetrahydrofolate reductase (MTHFR) as key enzymes involved in folate metabolism (Bai et al, 2019; Karatoprak et al, 2019; Padmanabhan et al, 2013) were gradually activated with the occurrence of ZGA in the development of mouse embryos (Appendix Fig S3D). Abnormal of genes related to folate metabolism enzyme results in decreased enzyme activity, which may cause blocked conversion of homocysteine (HCY) to methionine (MET). Folate also plays a critical role in the synthesis of S-adenosylmethionine (SAM) (Kim, 2000), which deficiency results decreased level of SAM, thereby resulting in abnormal methylation of Igf2 and other genes (Fig 5). Among them, we found that the knockout of Dppa2 and 4 down-regulates the expression level of Igf2 gene (Fig 5 and Appendix Fig S3D), which further causes the abnormality of signaling pathways regulated by Igf2, such as MAPK, PI3K-Akt and Ras signaling pathways (Aksamitiene et al, 2012; Ipsa et al, 2019). These pathways are also regulated by Dppa2/4, in agreement with the fact that the complex interactions between these signaling pathways are mainly involved in development and the alteration of cell fate. The lasted study from Gao’s group (Bi et al, 2020) reported that ALPPL2 interacts with IGF2BP1 to regulate human naïve pluripotency and overexpression of ALPPL2 can also led to significant activation of MAPK and ECM-receptor interaction. We can confirm that Alppl2 not only plays important roles in human naive pluripotency maintenance and establishment, but also was predicted as a potential master driver to regulate ZGA in signaling pathways.

## Conclusion

In our studies, we firstly profiled a genome-wide dynamic binding map of Dppa2 and Dppa4 among different cell types: MEFs, IPSCs and ESCs, the result implied that Dppa2/4 can further facilitate the strengthening of cell pluripotency, especially significant is cooperative effect of both. Compared with other TFs, Dppa2/4 are inclined to bind on GC-rich region of proximal promoter to activating majorities of signaling pathways. By comparing the binding characteristic of Dppa2 and 4 in three cells types, we observed that the binding abilities of Dppa2/4 in ESCs and IPSCs more strongly than in MEFs. Especially, the coordinately binding of Dppa2/4 on genes was superior to individual regulation, in which Dppa4 was more significant. Intriguingly, there was more substantial effect of Dppa4-bound on genes than Dppa2 in ESCs. Moreover, we found that Dppa2/4 can comprehensively active some signaling pathways associated with developmental reprogramming for promoting ZGA. Furthermore, we identified some directly targets of Dppa2 and 4. In which, Alppl2 was loss expression after knockdown of either Dppa2 or Dppa4, showing that Alppl2 is the direct downstream genes of these two factors and probably functions in ZGA. Strikingly, Alppl2 also involved in folate biosynthesis and can further regulates the complex interactions among key pathways associated with development. In conclusion, our studies hope to provide extensive new insights into the function and regulatory of Dppa2/4 in the process of early embryo development.

## Materials and Methods

### Dataset collection

The ChIP–seq data set of Dppa2/4 single- and double-overexpressed in MEFs, IPSCs and ESCs was downloaded from Gene Expression Omnibus (GEO) database under accession number GSE117171 (Hernandez et al, 2018). And the RNA-seq data of Dppa2/4 single-, double-knockout treatment (Dppa2-KO, Dppa4-KO and DKO) and control (WT) ESCs were also obtained in GEO database and GEO accession no.GSE120952 (Eckersley-Maslin et al, 2019). Moreover, the single-cell RNA-seq data of mouse embryos development (GSE70605) was reanalyzed in this study, which including two embryos types of somatic cell nuclear transfer (SCNT) embryos and in vitro fertilization (WT) embryos (Liu et al, 2016).

### ChIP-seq data processing

The ChIP-seq original fastq format data were controlled by FastQC software (http://www.bioinformatics.babraham.ac.uk/projects/fastqc/) to remove low-quality samples. Next the ChIP–seq reads were mapped to the mouse genome assembly Mm10 by using Hisat2 (Pertea et al, 2016) (version 2.1.0) short read alignment software with default parameters. Then we used MACS2 (De Iaco et al, 2019; Feng et al, 2011) (version 2.1.0) (with the parameters setting: macs2 callpeak -t $treatmentsam -c $controlsam -f SAM --keep-dup 1 -n $name -g 1.87e9 -B -q 0.01) to call binding peaks. Finally, using R package ChIPseeker (Yu et al, 2015) annotated with the position of the peaks in the genome, in which −2kb to 1kb of gene transcription start sites (TSS) were defined gene promoter. We also calculated the occupancy of Dppa2 and 4 in each peak as RPKM (reads per kb per million uniquely mapped reads) (Chen et al, 2016) for their binding signals.

### RNA-seq analysis

The RNA-seq original fastq format data of Dppa2/4 single-, double-knockout treatment (Dppa2-KO, Dppa4-KO and DKO) and control (WT) ESCs were controlled by FastQC software (http://www.bioinformatics.babraham.ac.uk/projects/fastqc/) to remove low-quality samples. The cleaned reads were mapped to Mm10 reference genome using (Pertea et al, 2016) Hisat2 (version 2.1.0) aligner for stranded and paired-end reads with default parameters. Reads count of gene expression was performed using HTseq-count (Python package). Next, transcriptome assembly was performed using Stringtie (Pertea et al, 2016; Pertea et al, 2015) (version 1.3.5) and Ballgown (Pertea et al, 2016) (R package), and expression levels of each genes were quantified with normalized FPKM (fragments per kilobase of exon model per million mapped reads) to eliminate the effects of sequencing depth and transcript length (Liu et al, 2016). Beside, differential expression analysis were conducted by R package DEseq2 (Love et al, 2014), for each comparison, genes with a Benjamini and Hochberg-adjusted P value (false discovery rate, FDR) < 0.05 and the absolute of Log2(fold change) > 1 were called differentially expressed (Liu et al, 2016).

### Functional enrichment and statistical analysis

Kyoto Encyclopedia of Genes and Genomes (KEGG) (Kanehisa et al, 2016) pathway enrichment analysis was performed based on the Database for Annotation, Visualization and Integrated Discovery (DAVID) Bioinformatics Resource (Huang da et al, 2009) (http://david.abcc.ncifcrf.gov/home.jsp). Statistical analyses were implemented with R (Aho) (version 3.6.0, http://www.r-project.org). Representative KEGG pathways were summarized in each gene cluster and P-values were marked to show the significance. The Pearson correlation coefficient was calculated using the ‘cor’ function with default parameters to estimate the correlation between genes (Ahlgren et al, 2003). The developmental data are represented as the average plus standard deviation of biological replicates (Mean+SD). Student’s t test was performed using the ‘t.test’ function with default parameters (Bandyopadhyay et al, 2014; Greenland et al, 2016).

### GC-content and CpG-Island analysis

To compute the GC-content of peaks bound by Dppa2/4, the peaks of both were done first by converting bed files to fasta with bedtools suite and then by using a home-made python script to count DNA bases. As for the CpG-Island annotations, which were downloaded from UCSC Genome Browser (CpG Islands).

### Motif-enrichment analysis

MEME-ChIP (Machanick & Bailey, 2011) was used to analyze Dppa2/4 binding motif with the parameter ‘-meme-mod zoops -meme-minw 4 -meme-maxw 10-meme-nmotifs 10 -meme-searchsize 100000 -dreme-e 0.05 -centrimo-score 5.0-centrimo-ethresh 10.0’, and sequences of top 1000 Dppa2/4 binding peaks were used as input. ‘vmatchPattern’ function from R package Biostrings was applied to determine the locations of Dppa2/4 binding motifs on Alppl2.

## Data visualization

In this study, data visualization was mainly achieved with R (version 3.6.0), including the R/Bioconductor (http://www.bioconductor.org) software packages. The heatmap, Upset plot and Venn plot were produced using R packet Pheatmap, UpsetR (Conway et al, 2017) and VennDiagram, respectively. The Integrative Genomics Viewer (IGV) (Thorvaldsdottir et al, 2013) was applied to visualize genome browser view. And the density graph, boxplot, PCA and so on were generated with the R packet ggplot2 (http://ggplot2.org/).

## Acknowledgments

The authors would like to thank the Prof. Shaorong Gao (Tongji University) and Prof. Yi Zhang (Harvard Medical School) for sharing their single-cell RNA-seq data of somatic cell nuclear transfer (SCNT) embryos in GEO database. The authors also thank Prof. Natalia B. Ivanova (Yale University) for sharing their Chip-seq data of Dppa2 and Dppa4 in GEO database; Prof. Wolf Reik (Babraham Institute) for sharing their RNA-seq data of Dppa2/4 single-, double-knockout treatment and control ESCs in GEO database. This work was supported by the National Nature Scientific Foundation of China (grants 61561036, 61702290, and 61861036); Program for Young Talents of Science and Technology in Universities of Inner Mongolia Autonomous Region (grant NJYT-18-B01); and the Fund for Excellent Young Scholars of Inner Mongolia (grant 2017JQ04).

## Author contributions

YZ conceived and designed the study. HL and CL did most of the bioinformatics analysis. JX and PL were helpful for materials collection. YZ and HL wrote the paper. All authors read and approved the manuscript.

