## Supplemental Figure S1-S3 for "Dppa2/4 promotes zygotic genome activation by binding to GC-rich region in signaling pathways"

**Table of Content**

Appendix Figure S1-S3 with Legends

**Appendix Figure S1**

**Comparison of chromosomes distribution of Dppa2 and 4 in MEFs, IPSCs and ESCs.**

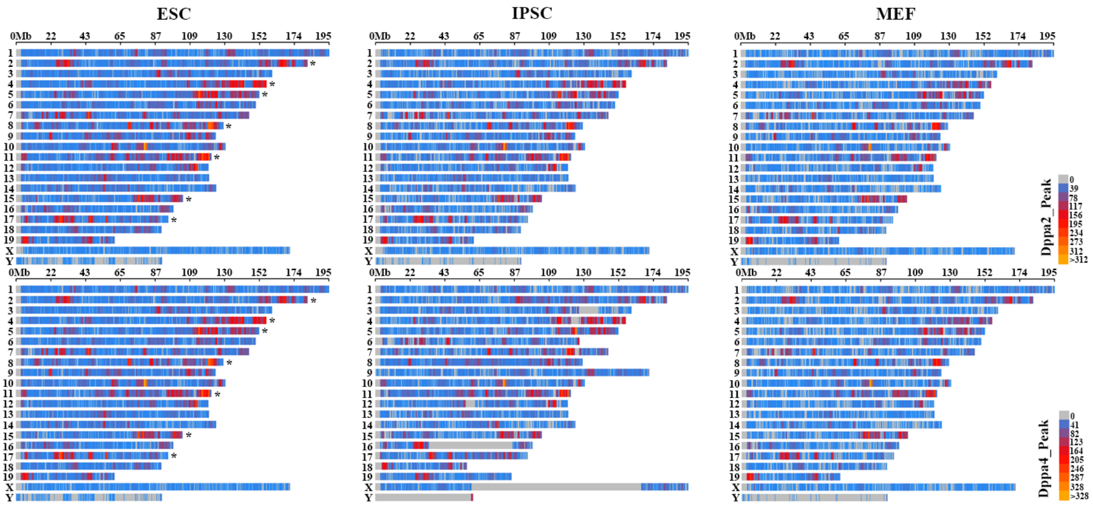


**Appendix Figure S2**

**The genome binding characteristics of Dppa2/4 in three different cell types**

(A) The percentage of target genes containing CpG Island in the target genes of Dppa2, Dppa4 binding onpromoter in all target genes containing CpG Island.

(B) Dppa2 and Dppa4-recognized DNA motifs identified by MEME-ChIP in MEFs and IPSCs, respectively.

(C) The target binding characteritics of Dppa2 and Dppa4 in the three different cell types.

(D) Significant enriched signaling pathways (adjPalve <= 0.5) and enriched target genes of Dppa2, Dppa4 and other TFs in ESCs (data reanalyzed from (Chronis *et al*, 2017) ).

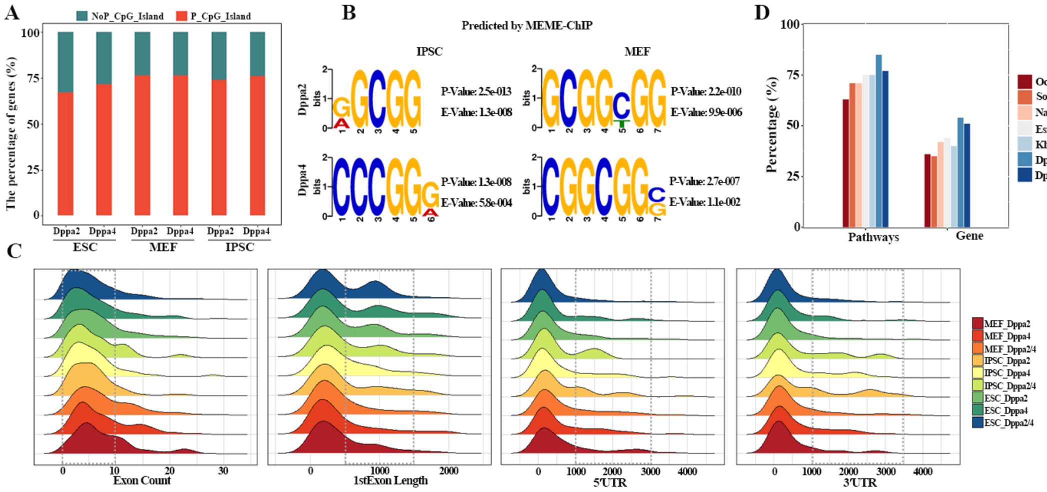


**Appendix Figure S3**

**Alppl2 is predicted as key factor functions in activating ZGA**

(A) The expression levels changes of ALPPL2、DPPA2、DPPA4 in the development of human embryos. Differences are statistically significant. (*) P-value < 0.05; (**) P-value < 0.01; (***) P-value < 0.001, t-test. Data are from Fan G et al. (Xue *et al*, 2013).

(B) The point plot shows the expression levels of ALPPL2、DPPA2、DPPA4 in the development of human embryos, both the point size and color represents normalized gene expression levels(Log2(FPKM+1)).

(C) Dynamic changes of representative genes in Figure 5 during the development mouse embryos.

(D) Genome browser views showing expression signals around Alppl2, Dppa2, Dppa4, Igf2 and Obox6 when Dppa2/Dppa4 single-, double-knockout (upper panel) and at each indicated mouse embryo development stage (bottom panel).

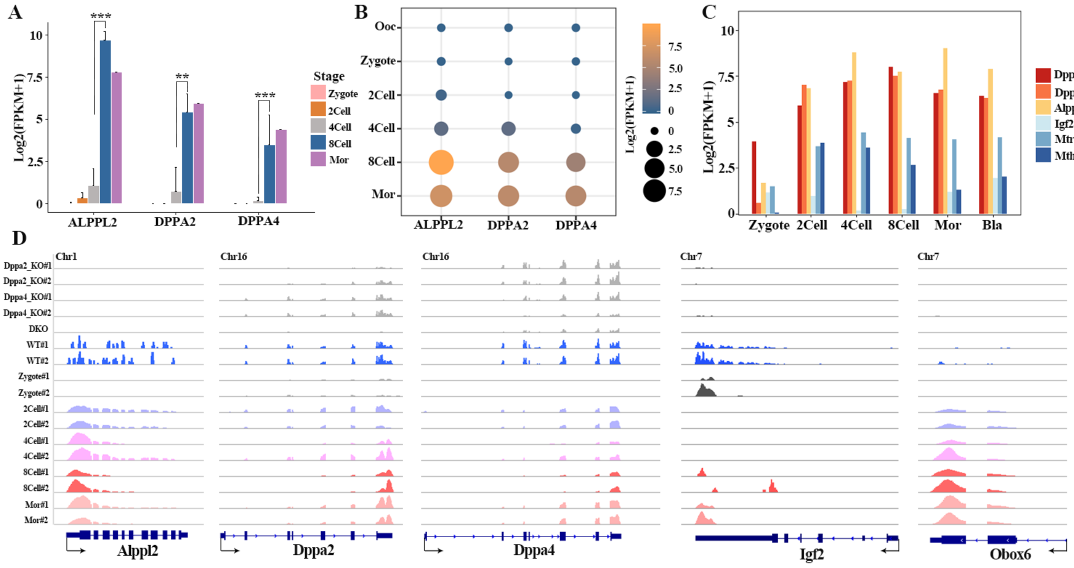
